# Digital pathology assessment of kidney glomerular filtration barrier ultrastructure in an animal model of podocytopathy

**DOI:** 10.1101/2024.06.14.599097

**Authors:** Aksel Laudon, Zhaoze Wang, Anqi Zou, Richa Sharma, Jiayi Ji, Connor Kim, Yingzhe Qian, Qin Ye, Hui Chen, Joel M Henderson, Chao Zhang, Vijaya B Kolachalama, Weining Lu

## Abstract

**Background:** Transmission electron microscopy (TEM) images can visualize kidney glomerular filtration barrier ultrastructure, including the glomerular basement membrane (GBM) and podocyte foot processes (PFP). Podocytopathy is associated with glomerular filtration barrier morphological changes observed experimentally and clinically by measuring GBM or PFP width. However, these measurements are currently performed manually. This limits research on podocytopathy disease mechanisms and therapeutics due to labor intensiveness and inter-operator variability.

**Methods:** We developed a deep learning-based digital pathology computational method to measure GBM and PFP width in TEM images from the kidneys of Integrin-Linked Kinase (ILK) podocyte-specific conditional knockout (cKO) mouse, an animal model of podocytopathy, compared to wild-type (WT) control mouse. We obtained TEM images from WT and ILK cKO littermate mice at 4 weeks old. Our automated method was composed of two stages: a U-Net model for GBM segmentation, followed by an image processing algorithm for GBM and PFP width measurement. We evaluated its performance with a 4-fold cross-validation study on WT and ILK cKO mouse kidney pairs.

**Results:** Mean (95% confidence interval) GBM segmentation accuracy, calculated as Jaccard index, was 0.73 (0.70-0.76) for WT and 0.85 (0.83-0.87) for ILK cKO TEM images. Automated and manual GBM width measurements were similar for both WT (p=0.49) and ILK cKO (p=0.06) specimens. While automated and manual PFP width measurements were similar for WT (p=0.89), they differed for ILK cKO (p<0.05) specimens. WT and ILK cKO specimens were morphologically distinguishable by manual GBM (p<0.05) and PFP (p<0.05) width measurements. This phenotypic difference was reflected in the automated GBM (p<0.05) more than PFP (p=0.06) widths.

**Conclusions:** These results suggest that certain automated measurements enabled via deep learning-based digital pathology tools could distinguish healthy kidneys from those with podocytopathy. Our proposed method provides high-throughput, objective morphological analysis and could facilitate podocytopathy research and translate into clinical diagnosis.

**Key points:**

- We leveraged U-Net architecture in an algorithm to measure the widths of glomerular basement membrane and podocyte foot processes.
- Deep learning-based automated measurement of glomerular filtration barrier morphology has promise in podocytopathy research and diagnosis.

## Introduction

Chronic kidney disease impacts over 13% of the global population.^1^ Proteinuric kidney diseases comprise a large portion of this massive health burden and often result in kidney failure.^2^ Podocytopathy is a proteinuric kidney disease caused by injury to the podocytes in the glomeruli.^3^ The podocyte is a critical part of the kidney’s glomerular filtration barrier that serves as a selectively permeable membrane to filter blood plasma while retaining blood cells and large serum proteins such as albumin.^1^ The glomerular filtration barrier also consists of the glomerular basement membrane (GBM) and fenestrated endothelium, which together perform kidney glomerular ultrafiltration function and prevent leaking of essential proteins into the urine (i.e., proteinuria).^2–4^ Podocytes are postmitotic cells with interdigitating, finger-like protrusions called foot processes that adhere to and compress the underlying GBM.^4^ Between podocyte foot processes (PFPs) are special cell-cell junctions with narrow gaps of about 40 nm called slit diaphragms (SD).^4^

In both research and clinical pathology settings, transmission electron microscopy (TEM) images of kidney tissues are commonly used to observe the glomerular filtration barrier ultrastructure and detect morphological changes (Figure 1A). TEM images provide a grayscale, cross-sectional visualization of the GBM, PFPs, and SDs, which are difficult to resolve under light microscopy (Figure 1B).^5^ The glomerular filtration barrier morphology can be quantified using TEM images by measuring the widths of the GBM and PFPs. GBM width is measured as the orthogonal distance between the basal cell membrane of the endothelium and the basal surface of the PFPs (Figure 1C).^6^ On the other hand, PFP width is measured as the distance between subsequent slit diaphragms parallel to the GBM (Figure 1C).^7^

**Figure 1.**
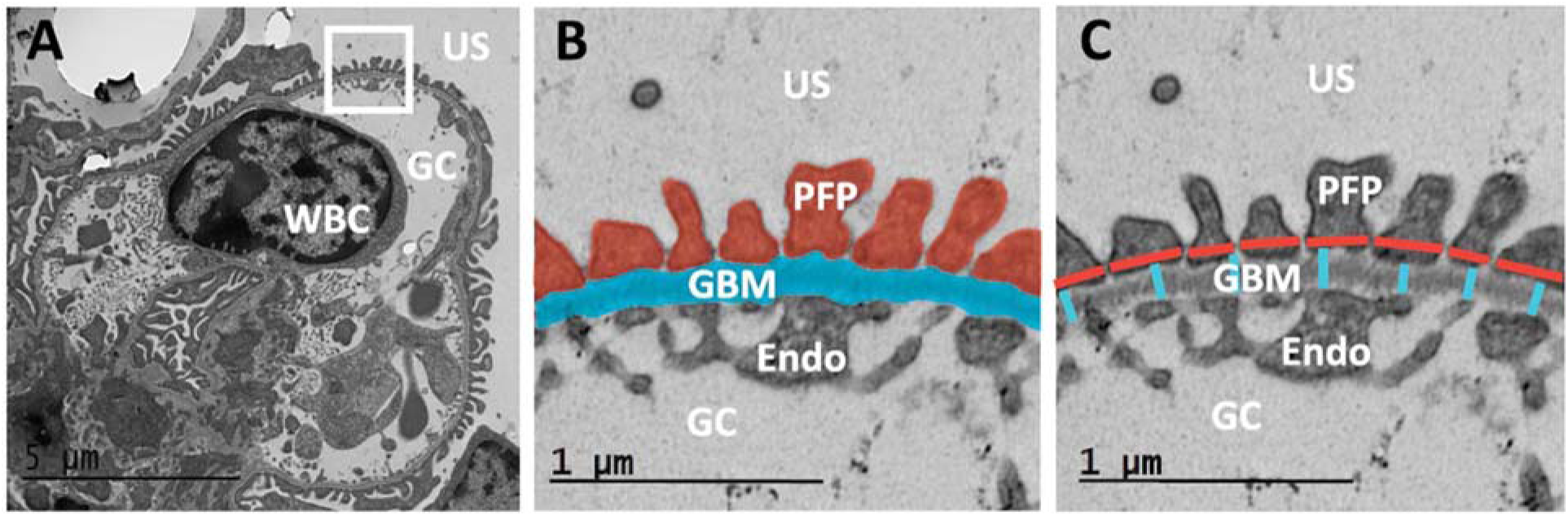
Illustration of kidney transmission electron microscopy (TEM) image ultrastructure and measurement. **(A)** Kidney TEM image of a wild-type mouse with a region of the glomerular filtration barrier in a white box. GC, glomerular capillary; US, urinary space; WBC, white blood cell. **(B)** The enlarged white box area in A shows the same glomerular filtration barrier region with highlighted podocyte foot processes (PFP, red) and glomerular basement membrane (GBM, blue). Endo, endothelial Cell. **(C)** Demonstration of manual measurement of PFP width (horizontal red lines) and GBM width (vertical blue lines).

These measurements taken from TEM images of patient kidney biopsies facilitate renal pathologists’ diagnosis and classification of glomerular diseases. An increase in PFP width, also called PFP effacement, has been associated with podocytopathies such as focal segmental glomerulosclerosis, minimal change disease, and membranous nephropathy (MN).^8,9^ Recently, restored PFP width was identified as an independent indicator of treatment outcome and remission in patients with proteinuric kidney disease such as lupus nephritis.^10^ Increased GBM width is often observed in podocytopathies such as MN^11^ and other proteinuric kidney diseases such as diabetic kidney disease^12,13^ and lupus nephritis,^5,14^ while decreased GBM width may suggest thin basement membrane nephropathy with type IV collagen mutations.^5,14^ These metrics are also used experimentally as image biomarkers for podocytopathy as they enable quantification of glomerular filtration barrier ultrastructural changes due to podocyte injury, genetic mutation, or therapeutic intervention.^7,15–17^

Despite the utility of GBM and PFP widths, efforts to fully automate their measurements in TEM images have been limited. Researchers and clinical renal pathologists continue to rely on manual measurement of TEM images, which is labor-intensive and limits the efficiency and accuracy of glomerular disease research and clinical diagnosis.^7,12,14,16,18–22^ In addition, manual measurement introduces significant inter-operator variation and bias,^6,19,22,23^ which affects reproducibility and can be addressed by automated measurement of kidney glomerular ultrastructure. Recent computation-aided tools measured either GBM or PFP width, though not both.^20,24^

In this study, we leveraged a deep learning framework to measure GBM and PFP widths in kidney TEM images from Integrin-Linked Kinase (ILK) podocyte-specific conditional knockout mice, an animal model of podocytopathy and severe proteinuria,^25,26^ and their wild-type littermate mice. Such frameworks have the potential to be translated for use with human kidney biopsy TEM images.

## Methods

### Animal Models

*ILK* podocyte-specific conditional knockout mice and their wild-type littermates were generated using the *ILK*^flox^ and NPHS2-*Cre* mice as previously described.^25^ The tail DNAs were isolated from 3-week-old mice produced from the breeding of ILK^flox/+^;NPHS2-Cre^+^ heterozygous mice. The ILK^flox/flox^;NPHS2-Cre^+^ homozygous mice (ILK cKO) and their wild-type (WT) littermates (ILK^+/+^;NPHS2-Cre^+^) were subsequently identified by polymerase chain reaction (PCR) genotyping methods, as previously described.^25^ At 4 weeks of age, WT and ILK cKO mice were euthanized, and their kidneys were dissected and fixed in Karnovsky’s fixative. TEM Images were obtained using a Joel JEM-1011 TEM instrument (JEOL USA Inc., USA) as described previously.^7,27^

### Manual Measurement of the GBM and PFP Width on TEM Images

Six to 12 TEM images at 12,000x magnification (3484-by-2672 pixels) were captured for each of the WT (n = 4) and ILK cKO (n = 4) mice, for a total of n = 40 TEM images from mice of each genotype. An expert operator manually measured GBM width on n = 30 images from WT and n = 23 from ILK cKO mice and PFP width on n = 23 images from WT and n = 28 images from ILK cKO mice using ImageJ software (version 1.51g; National Institutes of Health, USA). GBM width was measured between the basal surfaces of the endothelium and podocyte foot processes, orthogonal to the GBM contour,^5^ while PFP width was measured between subsequent slit diaphragms parallel to the GBM contour (Figure 1C).^7^ The results were then pasted on an Excel sheet and the harmonic mean for each image was calculated.

### U-Net Model and GBM Segmentation

The first step of our automated method was GBM semantic segmentation with a U-Net deep learning model.^28^ Ground truth GBM labels were produced from manually annotated TEM images with open-source QuPath software.^29^ The GBM was identified as described above, with the work divided among three operators.^6,21^ The resulting binary GBM labels classified each pixel as either GBM or background (Figure 2A). Each TEM image and its corresponding GBM label were shrunk row- and column-wise by a factor of two before a 100-pixel sliding window was applied to produce 8,586 overlapping 512-by-512 pixel label/image tile pairs from each whole image (Figure 2B). By utilizing overlapping crops and horizontal flips with a 50% probability, we augmented the size of our relatively small dataset. We trained and tested a U-Net model for GBM segmentation on this preprocessed dataset of TEM label/image tile pairs.^28^ This U-Net model is designed for image segmentation tasks, featuring 4 encoding blocks and 4 decoding blocks. It includes batch normalization after each convolution, ReLU activations, and the ‘same’ convolution mode to maintain spatial dimensions.

**Figure 2.**
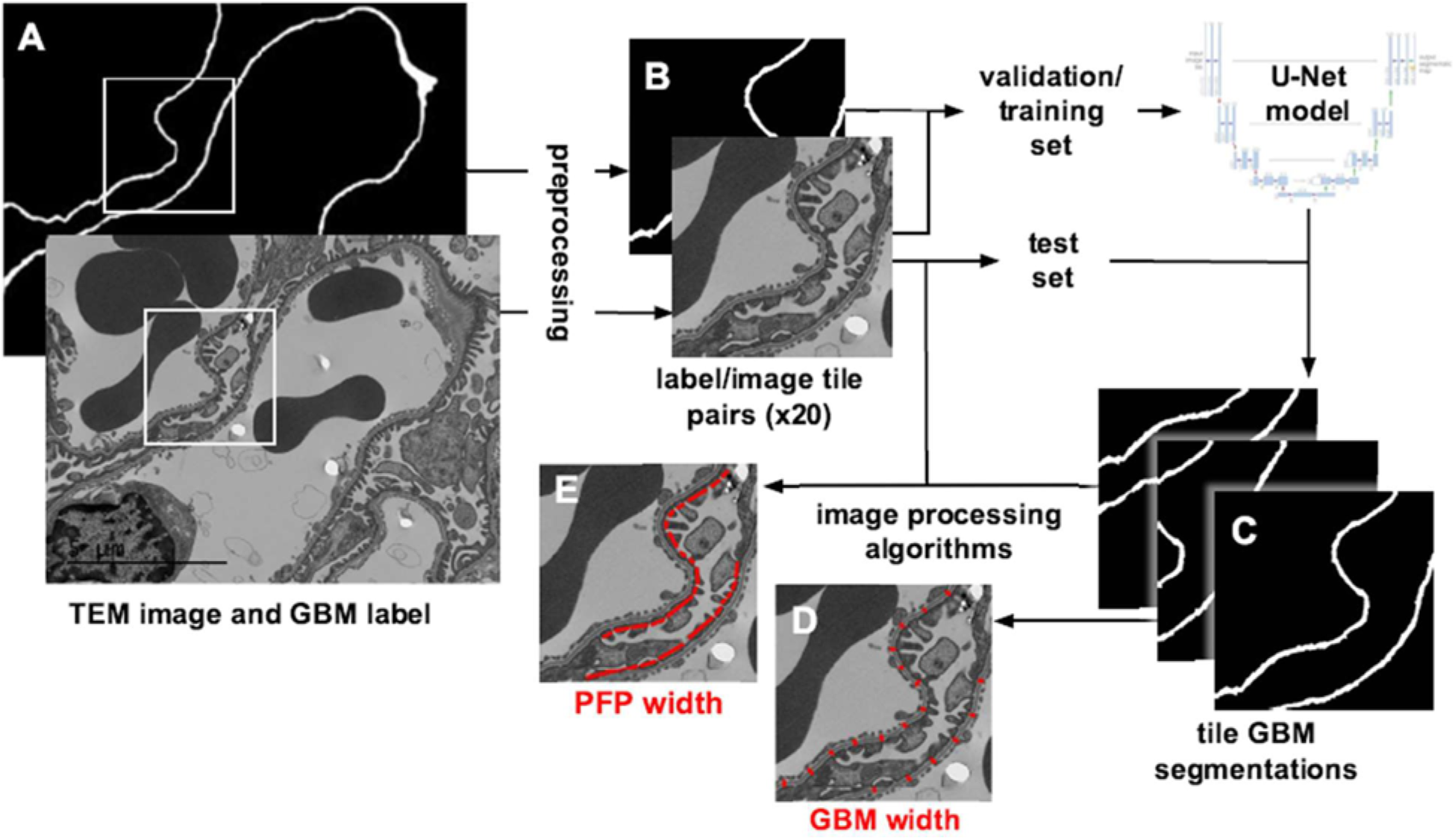
Block diagram of automated measurement method for glomerular basement membrane (GBM) and podocyte foot process (PFP) width. **(A)** Transmission electron microscopy (TEM) image and corresponding annotated GBM label. **(B)** Preprocessing results in 20 label/image tile pairs per label/image, which are sorted into validation/training and test sets, and the former are used to train a U-Net model for GBM segmentation. **(C)** The U-Net model is applied to test set TEM image tiles to generate GBM segmentation tiles. Resulting GBM segmentation tiles inform image processing algorithms for **(D)** GBM width, and along with image tiles, and for **(E)** PFP width.

During training, 25% of label/image tile pairs in the training set were randomly selected as the validation set to assess prediction accuracy after each training epoch and guide the update of model parameters. With each training epoch, our model seeks to minimize a dice loss (DL) function applied to the validation set predictions.

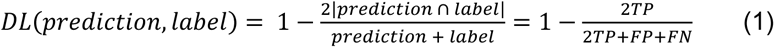

We selected 45 epochs as training length by plotting validation and training learning curves and observing the approximate epoch at which the losses were no longer reduced by additional training. A constant learning rate of 0.01 was selected with an open-source learning rate finder library by plotting validation loss as a function of the learning rate and choosing the approximate rate at which it was minimized. Additionally, we used a batch size of 2 and optimized the model using Stochastic Gradient Descent (SGD). U-Net models were trained with these optimized hyperparameters and saved after the final epoch. We then applied the U-Net to the test set TEM image tiles to perform GBM segmentation (Figure 2C), which was used to inform image processing algorithms for measuring GBM (Figure 2D) and PFP width (Figure 2E).

#### GBM Width Measurement Algorithm

The second stage was an image processing algorithm with the input of the predicted GBM segmentations. To measure mean GBM width, we first refined the segmentation results. We smoothed the GBM segmentation tiles with a narrow Gaussian kernel (standard deviation one pixel), applied Otsu-thresholding to generate a binary image, and removed small artifacts with areas less than 1,000 pixels, as these generally corresponded to false positive GBM. We then applied a skeletonizing algorithm to extract the center line of each GBM segment, as described in a previous publication (Figure 3A to 3B).^6^ Finally, the mean GBM width for each tile was calculated as its total GBM area divided by its total GBM central length:

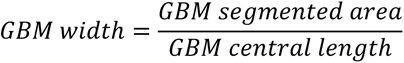

**Figure 3.**
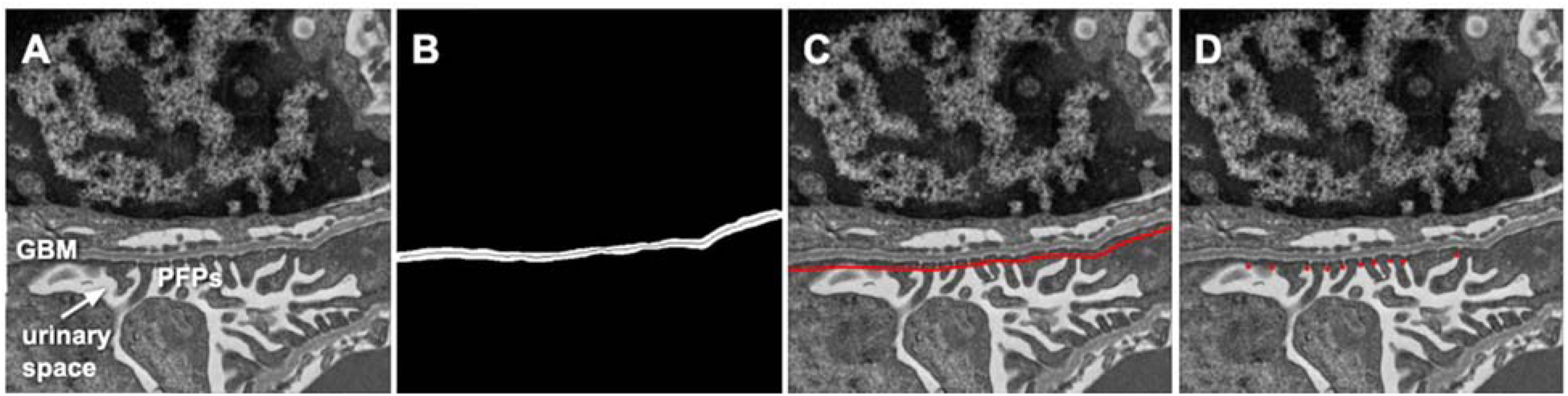
Glomerular basement membrane (GBM) and podocyte foot process (PFP) width measurement image processing algorithms. **(A)** Original transmission electron microscopy image tile with labelled GBM, PFPs, and urinary space. **(B)** Refined GBM segmentation results with one segment separated and fitted with a central line for GBM width calculation. **(C)** Urinary edge of smoothened GBM segmentation result. **(D)** Example identified slit diaphragms for PFP width calculation.

Our GBM width measurement algorithm borrows the approximation of GBM width as the quotient of the segmented area and central length,^30^ though we did not map the central length to a straight line, so we obtained a larger approximation. The mean GBM width for a given specimen was then calculated as the mean of its tiles’ GBM widths weighted by GBM area, then converted to length by a factor of 8.33 nm/pixel based on the scale bar.

#### PFP Width Measurement Algorithm

The predicted GBM segmentation tiles also guided a novel image processing algorithm for measuring PFP width. The segmentation tiles were first smoothed and widened by applying a Gaussian kernel (standard deviation 9 pixels) and then Otsu-thresholding. GBM tiles that had multiple distinct GBM segments with two edges were split into tiles with just one segment each. Tiles with discontinuous segments, with segments intersecting the tile borders at points other than the segment terminus, or without any segments, were discarded. This produced a refined dataset of GBM segmentation tiles cleanly traversed by the GBM segment. When these segmentation tiles with a widened, smoothened GBM were superimposed on the original TEM image tiles, the urinary side of GBM segments transected PFPs and slit diaphragms (Figure 3C). The TEM image pixel values were extracted from along both edges of this widened GBM segmentation. A Gaussian Mixture Model (GMM) classified the pixels along both edges into two classes by grayscale values corresponding to PFPs and slit diaphragms.^31^ The GMM assumed that each class’s values were normally distributed, which we confirmed was a rough approximation from histograms of several representative edges. The GMM predicted each class’s mean grayscale values and variances and applied these hyperparameters to classify the edge pixels into lighter versus darker classes. The urinary side was predicted as the edge with a higher proportion of pixels belonging to the darker class due to the dark staining of PFPs relative to the background. Distinguishing between the urinary and capillary edges of the GBM was a key step that required manual intervention in the most recent semi-automated PFP measurement effort.^23^

The pixel values from the urinary edge were plotted to identify the SDs. First, a smoothing Savitzky-Golay filter was applied to the one-dimensional data to reduce the impact of signal noise.^32^ The SDs were then identified from the smoothed edge values using a local maxima selection algorithm (Figure 3D). The minimum distance between SDs was set as 15 pixels, which corresponded to 125 nm and represented a conservative minimum PFP value from our expert manual measurement data. The normalized local maxima prominence threshold was set as the quotient of the standard deviation of the predicted GMM lighter/SD class and the edge mean pixel value. The mean PFP width for a given GBM segment tile was calculated as the quotient of the total urinary edge length and the number of identified SDs. To calculate the mean PFP width for a given specimen, the mean PFP width of its tiles was weighted according to the tile’s total urinary edge length and then converted to length by a factor of 8.33 nm/pixel.

#### Validation Study

We assessed our proposed automated measurement method using a 4-fold cross-validation study in which we trained 4 U-Net GBM segmentation models on 4 non-overlapping partitions of our preprocessed dataset of TEM image tiles and corresponding GBM labels (Figure 4). We first sorted our dataset specimens into 4 random pairs of one WT and one ILK cKO mouse. Four-fold cross-validation was conducted by (1) selecting one mouse pair as the test set and the remaining three pairs as the training set, (2) preprocessing the training and test set data, (3) training a U-Net GBM segmentation model, (4) predicting GBM segmentation of the test set, (5) using the predictions to measure GBM and PFP widths, (6) iterating the process 3 more times until each mouse pair served as the test set. In this manner, we approximated 75/25 training/test partitions and avoided overlap between training and test sets while maximizing the test data from our relatively small dataset. As a result, we obtained GBM segmentation for each TEM image and automated GBM and PFP width measurements for each mouse.

**Figure 4.**
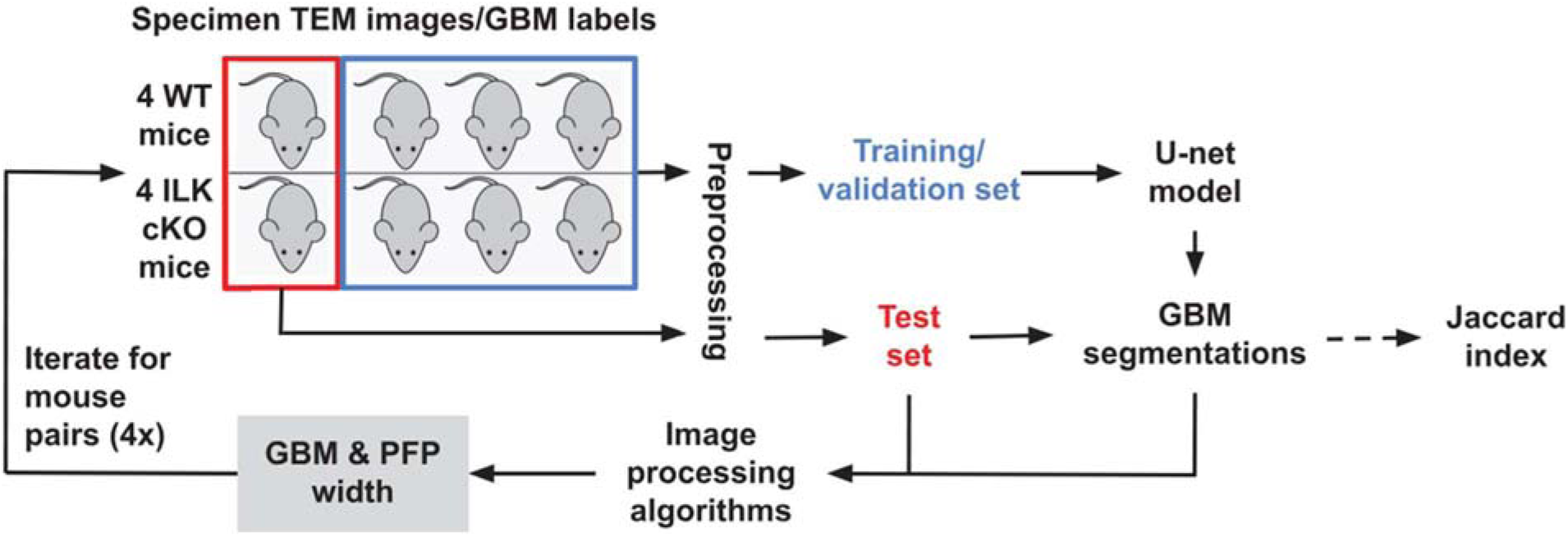
Four-fold cross-validation study of the proposed automated measurement method. TEM, transmission electron microscopy; GBM, glomerular basement membrane; WT, wild-type; ILK cKO; Integrin-Linked Kinase podocyte-specific conditional knockout; PFP, podocyte foot process.

GBM segmentation accuracy for each TEM image was reported as the Jaccard index (J), or intersection over union, often used to evaluate segmentation in biomedical images.^5,33^ We also used the Dice coefficient (D) to evaluate the accuracy. Both were calculated based on true positive (TP), false positive (FP), and false negative (FN) by comparing the predicted GBM segmentation tiles to their corresponding GBM label tiles.

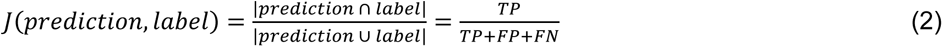

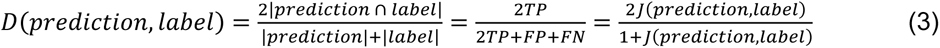

The Jaccard indices for each TEM image were averaged from its corresponding tiles. The resultant automated mean GBM or PFP width measurements for mouse specimens were compared with corresponding expert manual measurements for both WT and ILK cKO groups. Lastly, the manual and automated mean GBM and PFP width measurements for each mouse were compared between WT and ILK cKO groups.

#### Statistical Analysis

The mean Jaccard index values for both WT and ILK cKO TEM images were roughly normally distributed with a relatively large sample size (n = 40). Hence, we selected the parametric two-sample t-test to compare GBM segmentation accuracy between the two genotypes with an ɑ = 0.05 level of significance and reported the mean (95% confidence interval [CI]) image Jaccard index and Dice coefficient for both. The remaining statistical comparisons occurred between small (n = 4) groups with unknown distributions, so we selected the non-parametric Wilcoxon rank sum test with an ɑ = 0.05 level of significance. Data analyses were performed using RStudio (RStudio Inc., Boston, MA).

## Results

We evaluated our automated measurement of GBM and PFP width on our dataset of TEM images from 4 WT and 4 ILK cKO littermate mice with a 4-fold cross-validation study. GBM segmentation accuracy relative to the annotated labels for each TEM image is presented for WT and ILK cKO groups in terms of the Jaccard Index in Figure 5A. The resulting mean (95% CI) Jaccard indices for TEM images from WT and ILK cKO specimens were 0.73 (0.70-0.76) for WT and 0.85 (0.83-0.87) for ILK cKO specimen TEM images. This corresponds with Dice coefficients of 0.81 (0.79-0.84) and 0.88 (0.87-0.89), respectively. GBM segmentation accuracy was significantly greater for ILK cKO than WT images (95% CI, 0.07-0.14; p < 0.01, Figure 5A). TEM image tiles from WT and ILK cKO mouse kidneys are visualized in Figures 5B and 5D, respectively, alongside their corresponding predicted GBM segmentation in Figures 5C and 5E. On visual inspection, TEM image tiles with lower contrast and more irregular GBM morphology were generally less accurately segmented (Figures 5D and 5E)

**Figure 5.**
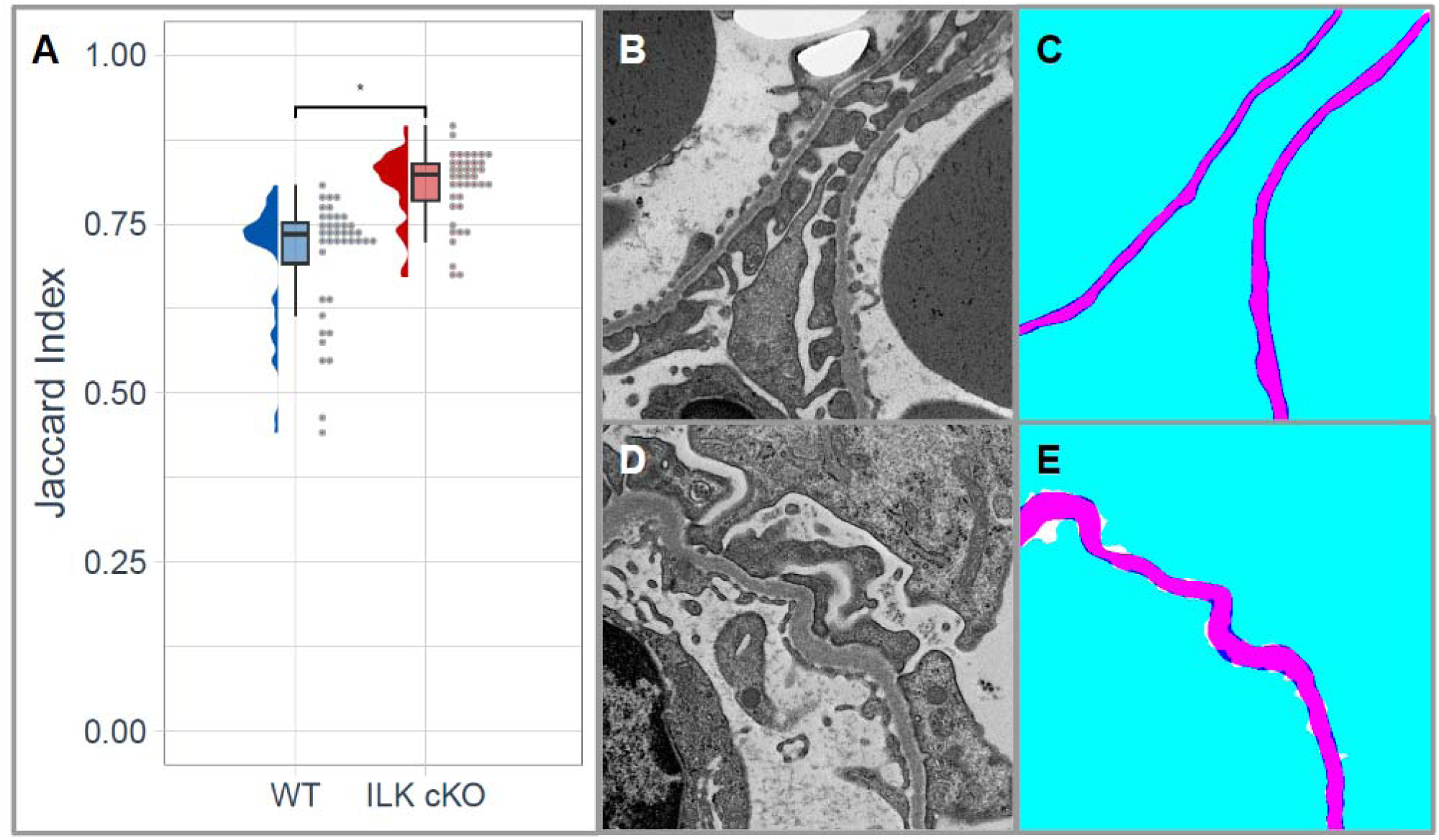
Accuracy of transmission electron microscopy (TEM) image glomerular basement membrane (GBM) segmentation with Jaccard index. **(A)** TEM image GBM segmentation accuracy for wild-type (WT) and Integrin-Linked Kinase podocyte-specific conditional knockout (ILK cKO) groups in terms of Jaccard index, where significance is denoted by p < 0.05 (*) in a two-sample t-test. **(B)** Representative TEM image tile from a WT specimen and **(C)** corresponding GBM segmentation results with true positive (pink), true negative (light blue), false negative (white), and false positive (dark blue). **(D)** Representative TEM image tile from an ILK cKO specimen and **(E)** corresponding GBM segmentation results with true positive (pink), true negative (light blue), false negative (white), and false positive (dark blue).

The automated measurements from the 4-fold cross-validation were compared with corresponding manual measurements of GBM (Figure 6A) and PFP width (Figure 6B) for both WT and ILK cKO groups. Manual and automated GBM width measurements were similar for WT (p = 0.49) and ILK cKO (p = 0.06) mice (Figure 6A). However, PFP width manual and automated measurements differed significantly for ILK cKO (p = 0.03) but not WT (p = 0.89) mice (Figure 6B).

**Figure 6.**
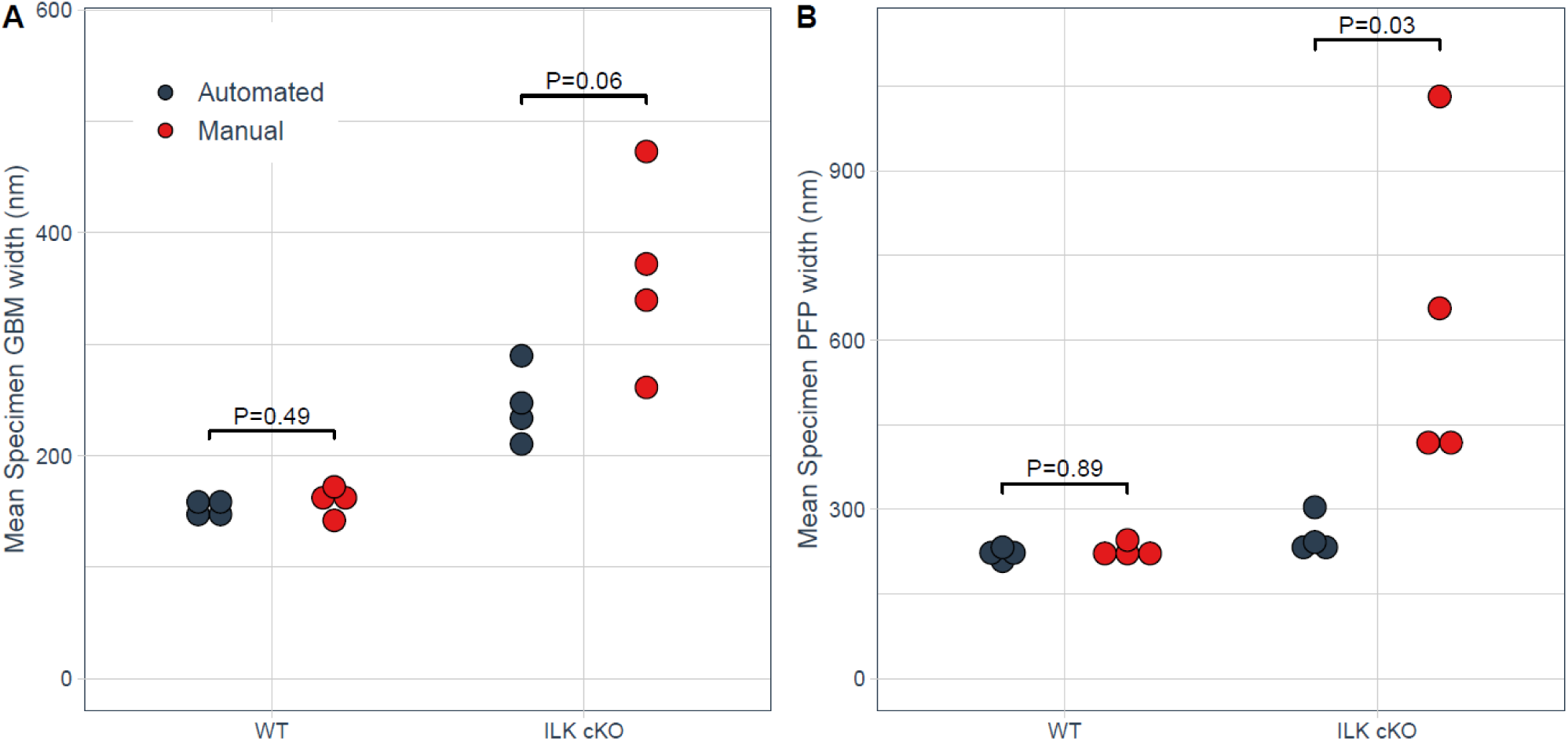
Comparison of automated versus corresponding manual measurements of glomerular basement membrane (GBM) and podocyte foot process (PFP) width in the same wild-type (WT) and Integrin-Linked Kinase podocyte-specific conditional knockout (ILK cKO) mouse kidneys. **(A)** Comparison of GBM width measured by automated versus manual method for WT and ILK cKO groups. **(B)** Comparison of PFP width measured by automated versus manual method for WT and ILK cKO. The exact p values are shown, which are calculated from the Wilcoxon rank sum test with an ɑ = 0.05 level of significance.

We also compared the GBM and PFP width between WT and ILK cKO groups with both automated (Figure 7A) and manual measurements (Figure 7B). Manual measurements differed significantly between genotype groups for both GBM (p = 0.03) and PFP width (p = 0.03) (Figure 7B). Similarly, automated GBM width measurements for WT mice differed significantly from ILK cKO mice (p = 0.03) (Figure 7A), although automated PFP width measurements differed less substantially (p = 0.06) (Figure 7A), which is consistent with the previously reported pathology finding in 4-week young ILK cKO mice.^26^

**Figure 7.**
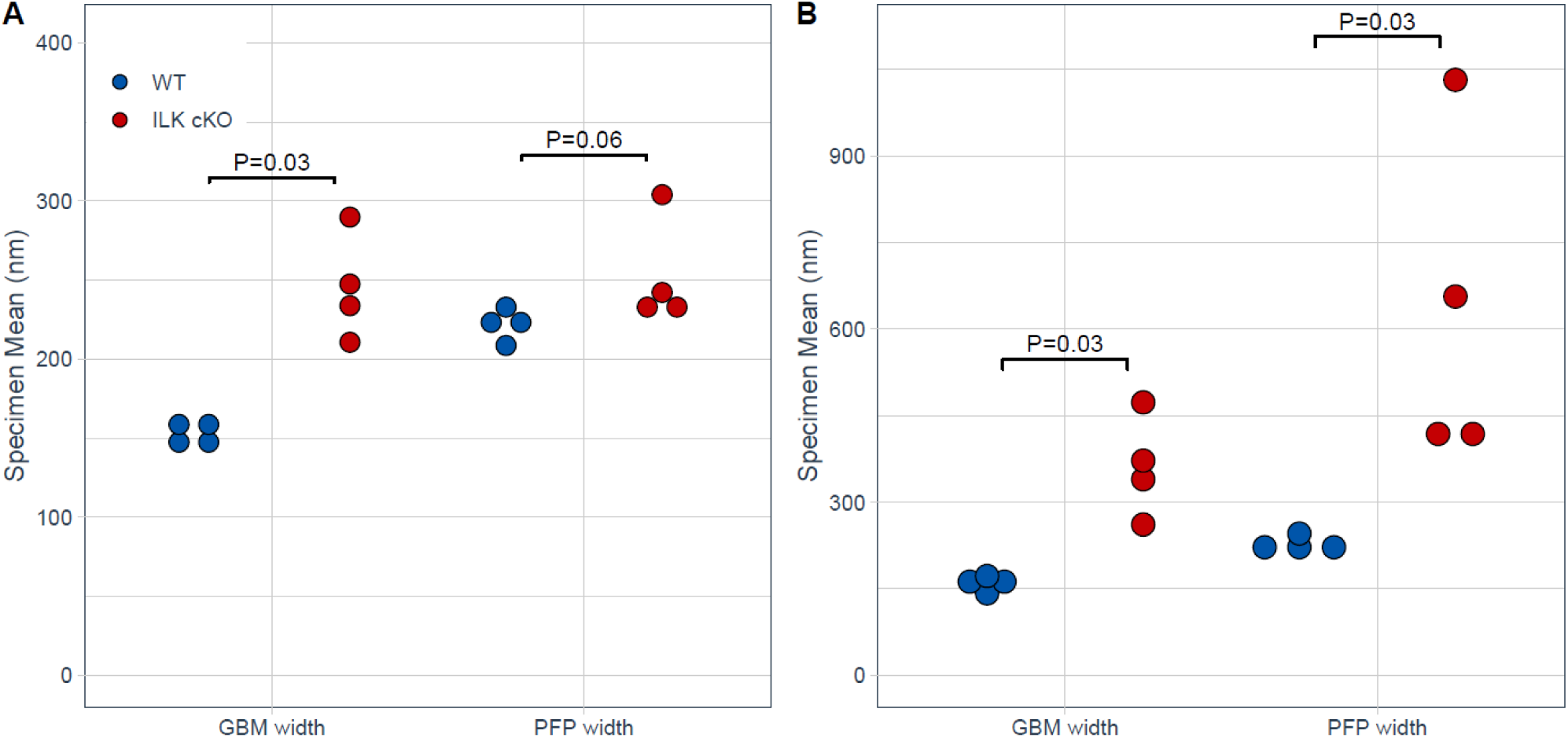
Distinction of subtle differences in glomerular basement membrane (GBM) width and podocyte foot process (PFP) width in 4-week-old wild-type (WT) versus Integrin-Linked Kinase podocyte-specific conditional knockout (ILK cKO) mouse kidneys measured by automated and manual methods. **(A)** Automated measurements of GBM width and PFP width in 4-week-old WT versus ILK cKO littermate kidneys. **(B)** Manual measurements of glomerular basement membrane GBM width and PFP width in 4-week-old WT versus ILK cKO littermate kidneys. The exact p values are shown, which are calculated from the Wilcoxon rank sum test with an ɑ = 0.05 level of significance.

## Discussion

In this work, we developed a method of automating GBM and PFP width measurement in kidney glomerular TEM images using a U-Net framework for GBM identification, followed by image processing measurement algorithms. We evaluated the segmentation and measurement performance through a 4-fold cross-validation study. Our automated GBM and PFP width measurements were similar to corresponding manual measurements for WT but not ILK cKO specimens. However, the manual measurements may be biased as ground truth, especially with a single operator. Instead, we emphasize that our approach may still serve to recognize pathological tissue. Our automated method distinguished early pathological changes in the GBM width more so than PFP width in young 4-week-old ILK cKO mouse kidneys. This result was consistent with findings in a previously published report of GBM derangements in ILK cKO mice, including increased GBM thickness, without a significant change in PFP width at 4 weeks.^26^ Our approach thus represents an advancement in quantifying glomerular TEM images, enhancing the speed and accuracy of analysis compared to current manual and semi-automated methods. Furthermore, our method demonstrates the potential for deep learning to augment podocytopathy experimental analysis and clinical diagnosis, which currently rely on the traditional paradigm of manual measurement.

Semi-automated methods have been piloted for GBM^6,12,30^ and PFP measurement^18,23^ but require operator intervention for each image. Automated measurement of PFP width has been developed for other imaging modalities, such as three-dimensional structured illumination microscopy (3D-SIM)^34^ and confocal and stimulated emission depletion (STED) super-resolution microscopy.^19^ However, unlike TEM, 3D-SIM and STED super-resolution microscopy are not commonly used in clinical pathology diagnosis and do not permit GBM width measurement. Recently, an automated method for measuring GBM width in TEM images was reported.^20^ However, this method did not include PFP width measurement and has not been validated.

Similarly, an automated method for PFP width measurement in kidney biopsy TEM images was also recently published.^24^ Compared to these two, our new “digital pathology” automated method can measure the GBM and PFP widths simultaneously.

We are among the first to apply a U-Net framework to GBM segmentation, joining Wang et al.,^20^ Yang *et al.,*^35^ and Smerkous *et al.*^24^ Deep learning models such as U-Net have become favored for various biomedical segmentation tasks due to robust learning from a small number of labeled images.^27,36^ The GBM may be particularly difficult to identify in TEM images given its variable morphology and poor image contrast.^36^ Unlike prior partially automated approaches, our U-Net represents fully automated GBM and PFP segmentation in TEM images.^5,6,12,18^ In addition, with Jaccard indices of 0.73 (0.70-0.76) for WT and 0.85 (0.83-0.87) for ILK cKO mice, our segmentation accuracy exceeded our initial goal of 0.5 and the 0.64 achieved by Cao *et al*’s random forest stacks-based machine learning.^5^ Although GBM morphology appeared more variable and indistinguishable in ILK cKO specimens, segmentation accuracy was significantly higher for this group. This finding may be explained by the mathematical tendency of the Jaccard index to favor images with higher counts of true positive pixels, given the wider mean GBM width of ILK cKO specimens. One disadvantage of our current segmentation framework was its reliance on GBM manual annotation for learning, which was itself labor-intensive.^20,36^ However, Wang et al.^20^ and Lin *et al.*^36^ recently superseded this limitation by pioneering a self-supervised approach for GBM segmentation that enables learning largely from unlabeled TEM images.

High-fidelity GBM segmentation guided our subsequent measurement algorithms. To the best of our knowledge, we are the first to fully automate both PFP width and GBM width measurement simultaneously in kidney TEM images. Similar to Smerkous *et al.*,^24^ a primary characteristic of our PFP width measurement algorithm that improves upon the approach of Vargas *et al*.^23^ to enable full automation is its ability to predict the urinary versus the capillary sides of the GBM, which we performed using GMM classification. However, slit diaphragm identification remains a key area of potential improvement, as visual inspection of algorithm-identified slit diaphragms suggested frequent misidentification.

An important limitation of this study was the relatively small animal dataset, which included only one type of disease model. With only 4 specimens per genotype, our ability to draw strong conclusions about the accuracy of our automated measurements and their effectiveness in distinguishing pathological tissue was limited. Additionally, the small TEM image dataset (n = 80) restricted the robustness of our trained U-Net models, even with data augmentation techniques. Future studies may incorporate larger image datasets with more diverse pathologies, other animal models such as rats, and human kidney biopsies. Given the morphological similarity between human kidneys and those of mice, we anticipate that our method could be generalized with appropriate parameter scaling. Further research might also explore semi-supervised^20,36^ or reinforcement learning approaches to reduce the burden of labeling larger image datasets. Our long-term goal is to develop a web-based tool that researchers and pathologists can use to input TEM images and obtain automated GBM and PFP width measurements, thereby facilitating clinical diagnosis and research.

## Acknowledgments

We thank Dr. Diane Joseph-McCarthy and Dr. Darren Roblyer for their support on the Boston University Biomedical Engineering Senior Design Course. This work was supported in part by the National Institutes of Health grant R01-DK133940 (WL), R43-DK134273, R01-HL159620, R21-CA253498, and RF1-AG062109 (VBK), the DOD grant E01HT9425-23-1-1058 (WL), a grant from Boston University Biomedical Innovation Technologies Affinity Research Collaboratives (BIT-ARC), a Boston University Clinical & Translational Science Institute Integrated Pilot Grant (CZ, WL), and funding support from Boston University Undergraduate Research Opportunities Program (AL). This publication is also supported in part by the National Center for Advancing Translational Sciences, National Institutes of Health, through Boston University Clinical & Translational Science Institute Grant Number 1UL1TR001430. Its contents are solely the responsibility of the authors and do not necessarily represent the official views of the NIH.

## Author contributions

AL and WL conceived and designed the project. ZW, AL, and AZ developed the machine learning model. AL, CK, YQ, JJ, and QY developed the image processing algorithms and annotated the GBM labels. RS and HC generated ILK cKO mice and kidney TEM images. AL and CZ provided statistical analysis. CZ, VBK, JMH, and WL provided reagents/materials/analysis tools. AL, AZ, CZ, and WL wrote the manuscript. All authors read, edited, and approved the final manuscript.

## Data Sharing Statement

The segmentation model and code are publicly available at https://github.com/CZCBLab/GBM_UNET. The data set used and/or analyzed during the current study is available upon request.

